# Genome-Wide Associations within Diverse Wild Apple Germplasm for Postharvest Blue Mold Resistance to *Penicillium expansum*

**DOI:** 10.1101/2024.12.30.629434

**Authors:** Lauren Whitt, John S. Bennett, Tamara D. Collum, Breyn Evans, Doug Raines, Ben Gutierrez, Wojciech J. Janisiewicz, Wayne M. Jurick, Christopher Gottschalk

**Affiliations:** Appalachian Fruit Research Station, USDA-ARS, Kearneysville, WV; Plant Genetic Resource Unit, USDA-ARS, Geneva, NY; Food Quality Lab, USDA-ARS, Beltsville, MD

**Keywords:** blue mold resistance, *Malus sylvestris*, *Malus sieversii*, DNA markers, GWAS, postharvest decay, patulin

## Abstract

Post-harvest disease caused by the blue mold fungus, *Penicillium expansum*, accounts for a substantial proportion of economic losses in United States apple industry. Multiple modes of entry in the apple supply chain, plus emerging fungicide resistance, limit the current and long-term viability of using chemical controls alone. Previous phenotypic screens of *Malus* accessions in the USDA-ARS apple germplasm have identified varying levels of blue mold disease resistance in some wild apple accessions and hybrids. These wild apple species contain reservoirs of genetic resistance that can be integrated into apple breeding programs to complement the previously identified *qM-Pe3.1* marker from *M. sieversii*. We sought to identify these novel loci by combining historical phenotypes of the USDA-ARS wild apple germplasm with low-pass genomic sequencing to perform association mapping. Multi-locus mixed models identified five single nucleotide polymorphisms (SNPs) significantly associated with reduction of post-harvest rot under high concentration of *P. expansum* inoculum, and one SNP associated under low inoculum concentration. Within a 25,000 base pair window of these SNPs, we found candidate genes encoding proteins with known pathogen immune response and defense roles, such as a Cobra-like 7, flavin monooxygenase, LRR receptors, PR5-like receptor kinase, and a putative resistance protein RGA3. We present these loci as targets for identifying accessions with beneficial alleles that can be targeted for fine mapping and use in *Malus* breeding programs to achieve *M. domestica* lines with natural post-harvest rot resistance.

## 1. Introduction

Apple (*Malus × domestica* Borkh.) is a highly valued tree fruit crop in the U.S. with farmgate values exceeding $3.05B (Koong et al., 2023). The vast majority of that $3.05BM is derived from fresh market sales (*e.g.* store-bought and direct-to-consumer sales) but about ∼14% is derived from alternatives such as juice, sliced, and other processed apple products (Koong et al., 2023). Although the majority of fruit produced enters the supply chain destined for either of these two avenues, a considerable amount is lost due to various pre- and post-harvest issues. Pre-harvest problems include premature fruit drop, appearance issues (e.g. lack of peel color or russeting), diseases such as apple scab, fire blight and bitter rot, and fruit size and shape (Arseneault and Cline, 2016; Kalcsits et al., 2019; Khodadadi et al., 2023; Musacchi and Serra, 2018; Van der Zwet et al., 2012). Post-harvest challenges include physiological defects and disorders associated with storage, mechanical damage to fruit during handling, sorting, bagging, bin-filling, and rot (Janisiewicz and Jurick 2017; Johnston et al. 2002; Jurick II et al. 2023; Mditshwa et al. 2018; Morales et al. 2010; Sidhu et al. 2023; Watkins 2017; Zhang et al. 2018). Of these challenges, post-harvest disease management is a major limitation and blue mold caused by *Penicillium expansum* is one of the most damaging (Argenta et al., 2021; Głos et al., 2022; Luciano-Rosario et al., 2020). Estimated losses from blue mold infection range from 1-5% on fungicide-treated fruit, thus impacts to the farmgate value of apples could be in excess of $18M (Luciano-Rosario et al., 2020).

*P. expansum* is a necrotrophic fungus that enters through lenticels, punctures, cracks, and other injuries on the apple peel (Errampalli, 2004; Luciano-Rosario et al., 2020; Rosenberger et al., 2006; Wenneker and Thomma, 2020). Upon entry into the fruit cortex, the fungi produces cell wall-degrading enzymes, acidifying factors such as PacC and organic acids, necrosis and ethylene-inducing proteins, and mycotoxins including patulin (Luciano-Rosario et al., 2020; Morales et al., 2010). The best method to control *P. expansum* is by application of fungicides by pre-harvest sprays and post-harvest fogs or dips (Edney, 1970; Little et al., 1980; Luciano-Rosario et al., 2020; Xiao and Boal, 2009, 2009). Moreover, biological controls have been tested, released, and approved for use (Janisiewicz and Jeffers, 1997). However, fungicide resistant strains have been found which creates needs for new means of control (Amiri et al., 2017; Jurick II et al., 2018). Biological controls have been developed tested, released, and approved for commercial use but it works best in combination with other alternatives to synthetic fungicides (Janisiewicz and Jeffers, 1997; Janisiewicz and Korsten, 2002).

Most durable disease control in crops is based on host-resistance. Cultivated varieties of apple completely lack host resistance to *P. expansum* (Luciano-Rosario et al., 2020; Spotts et al., 1999). However, resistance has been documented in progenitor species such as *M. sieversii* and *M. sylvestris* (Janisiewicz et al., 2008; Jurick II et al., 2011; Norelli et al., 2017). Initial efforts to map resistance loci were conducted using a population of ‘Gala’ × *M. sieversii* (Norelli et al., 2017). These efforts led to the development of a functional marker on chromosome 3, *qM-Pe3.1,* which has already been deployed in a pre-breeding program (Luo et al., 2020; Norelli et al., 2017). Although a significant achievement, this marker was developed from one *M. sieversii* accession catalogued as resistant (Jurick II et al., 2011). In comparison, different *M. sieversii* accessions and other *Malus* species have also exhibited high levels of resistance (Janisiewicz et al., 2016; Jurick II et al., 2011). These novel genetic resources could offer new resistant mechanisms and/or increased durability over *qM-Pe3.1* (Janisiewicz et al., 2016). However, no genetic analyses have attempted to elucidate the underlying resistance in this diverse *Malus* germplasm material.

In this investigation, we leveraged an opportunity to utilize this vast historical phenotypic information from Jurick II et al. (2011) to investigate these novel mechanisms and/or regions contributing to *P. expansum* host-resistance. Our approach utilized an emerging, cost-effective method for genotyping using whole genome resequencing at reduced coverages (∼3x) (Korani et al., 2021). We then applied a multi-locus, mixed model genome-wide association study (MLMM GWAS) to identify SNPs that associate with the quantitative phenotypic measurements from the Jurick II et al. (2011) study. With this approach, we aimed to elucidate new genetic resources for development of durable blue mold resistance for apple breeding programs from historical phenotypic information. We envision that this work will not only move the science of host resistance forward in a significant manner through its fundamental contributions but will also impact the pome fruit industry to supply them with a major source of resistance that can be introduced to commercial apple cultivars to help fight postharvest decay that is currently lacking.

## 2. Materials & Methods

### 2.1 Plant Material

In the Jurick II et al. (2011) manuscript, they evaluated a total of 452 accessions in the USDA apple collection located in Geneva, NY and accessions located at the USDA-ARS Appalachian Fruit Research Station (AFRS) in Kearneysville, WV. We extracted the accession numbers for all 452 accessions and cross referenced them with the current inventory lists of the USDA apple collection and USDA-ARS-AFRS. A total of 106 non-redundant accessions were found remaining of the 452 that were initially phenotyped in 2011. Of the 106 accessions remaining, six were located at AFRS while the other 100 are at the Geneva, NY collection. Fresh leaves were collected in the fall of 2022 from both locations and lyophilized on a VirTis Sentry 2.0 (SP Scientific; Warminster, PA) inside coin envelopes. Freeze-dried leaves were then stored at room temperature until DNA was extracted within two weeks of collecting.

### 2.2 Genomic DNA Extraction

Each accession had at least 0.5 grams of freeze-dried tissue ground in a mortar and pestle until a fine powder was produced. The powder was transferred to a 1.5 mL microfuge tube and 0.6 mL of 55°C D-TAB buffer was added to the tube and vortexed thoroughly for 30 seconds. The sample was then incubated at 55°C in a water bath for 30 minutes with occasional mixing. Following incubation, 600 µL of chloroform was added to the sample and vortexed for 30 seconds until thoroughly mixed. The sample was centrifuged at 12,000 rpm for 10 minutes. The supernatant was removed and placed into a fresh microfuge tube. The chloroform addition, mixing, and centrifugation was repeated a second time. Following collection of the supernatant, 600 µL of isopropanol was added and vortexed for 15 seconds and placed on ice for 15-30 minutes. The chilled mixture was centrifuged at 10,000 rpm for 10 minutes to pellet the DNA. The pellet was washed with 70% ethanol once and air-dried. DNA was then resuspended in TE buffer. The DNA was then cleaned and concentrated using Zymo Genomic DNA kit (Irvine, CA) following the manufacturer’s protocol. DNA quantification and purification quality was assessed using a Qubit fluorometer and Nanodrop spectrophotometer (ThermoFisher Scientific, Pittsburg, PA).

### 2.3 Genome Sequencing and Variant Calling

The extracted gDNA was sent to HudsonAlpha Institute for Biotechnology (Huntsville, MS) for library preparation and sequencing through the Khufu service (https://www.hudsonalpha.org/khufudata/). Khufu is a low-pass sequencing method that utilizes Illumina sequencing platforms (Korani et al., 2021). The resulting sequencing reads were demultiplexed, filtered, and processed using Khufu’s internal pipeline and mapped to a reference genome. For submission, the DNA samples were split across two 96-well sample plates. For the mapping-based variant calling, we utilized the phased Honeycrisp v1.0 hap 1A genome as the reference using BWA-mem (Khan et al., 2022; Li and Durbin, 2009). The genome was retrieved from the Genome Database for Rosaceae (GDR; Jung et al. 2019). Variant calls, imputation, and haplotype map preparation was performed using Khufu’s internal pipelines (https://www.hudsonalpha.org/khufudata/) (Korani et al., 2021). Single nucleotide polymorphisms (SNPs) missing calls from more than 75% of accessions and accessions missing more than 90% of SNPs were removed before imputation. Imputed genotypes were filtered to remove SNPs with a minor allele frequency (MAF) less than 5%.

### 2.4 Blue-mold resistance phenotyping

To review the complete methods for the phenotyping refer to Jurick II et al. (2011). In brief, fruit was harvested from each accession when iodine starch test values were >3 but <6 (Smith et al. 1979). Following harvesting, 20 fruit per accession were wounded with 6-penny nail to a depth of 3 mm. Following wounding the fruit was inoculated with a 50 μl conidial suspension of *Penicillium expansum* (MD-8). The wounding and inoculation experiment was conducted at two conidial suspension concentrations of 1 x 10^3^ (low) and 1 x 10^4^ (high) conidia/mL. Diameter of the lesions were then measured using a micrometer and recorded for each individual fruit. These experiments were conducted across three years between 2007 and 2009. The lesions sizes per year were used as the quantitative data for the GWAS. For the categorical classifications used in the GWAS, we relied on the classification results published in Jurick II et al. (2011). Susceptible accessions exhibited decay at both inoculation concentrations. Moderately resistant accessions exhibited decay at the 1 x 10^4^ concentration but not at the 1 x 10^3^ concentration. Resistant accessions exhibited no decay at either concentration.

The phenotypes for GWAS were the model estimated means of each accession within (*diameter*∼ *accession*) and across treatments (*diameter* ∼ *treatment* + *accession*). We used generalized linear models with binomial and negative binomial distributions for the binary and quantitative data, respectively. Resulting phenotypes with non-normal distributions were Box-Cox transformed to meet assumptions of the GWAS model.

### 2.5 Historical data curation

As is the case with most historic data, some measurements were missing or suspected to be swapped between accessions. Accessions that we could not confidently assign binary or quantitative measurements were removed from the study (PI613993, PI613998, PI59004, PI633801, and PI594107). Nine susceptible accessions and two moderately resistant accessions were missing the quantitative lesion measurements, but still had complete binary lesion data. We estimated quantitative measurements for these eleven accessions using mean imputation. For each of these accessions, we generated an estimated mean for their quantitative data by taking the average lesion diameter of all accessions in the same disease severity category (susceptible or moderate) and treatment (low/high) categories with similar 0/1 distributions in the binary data. For example, a susceptible accession missing all quantitative measurement has 15 binary measurements with a phenotype measurement of “1” under the low inoculum treatment (i.e. 15/20 samples have blue mold lesions). The equivalent mean of that susceptible accession’s low treatment is then set to the 0.75 (15/20) quantile of the low treatment measurements of all susceptible accessions with complete quantitative measurements. While mean imputation for missing phenotype data can reduce statistical power, and therefore GWAS signal, by shrinking values towards the mean (Poyatos et al. 2018; Gelman and Hill 2006), we wanted to retain as many individuals as possible for GWAS as the population already contained a relatively low number of accessions.

### 2.6 Genome-wide association studies

We conducted multi-locus mixed model (MLMM) GWAS for phenotypes under low and high *P. expansum* inoculum treatments in R (R Core Team, 2024) using the ‘mlmm’ package (Segura et al., 2012). The ‘mlmm’ package performs a stepwise, mixed-model regression, where SNPs are added as cofactors to the model based off their association with the phenotype. The maximum model is reached when all the variance due to estimated heritability is explained by the cofactors in the final step, or a maximum number of steps set by the user (default = 10) is reached. To control for false positives, the package includes two model selection criteria: extended Bayesian information criterion (extBIC) and multiple Bonferroni criterion (mBonf). To control for population structure, the model requires a kinship matrix, which we generated using the Astle-Balding method (Astle and Balding, 2009) from the ‘ionomicsUtils’ R package (v1.0 https://github.com/gziegler/ionomicsUtils; Wimmer et al. 2012). We also used the ‘mlmm’ option to include principal components as fixed covariates to the MLMM. We conducted a principal component analysis (PCA) with the EIGENstrat method (Price et al., 2006) implemented in the ‘ionomicsUtils’ package (v1.0 https://github.com/gziegler/ionomicsUtils). We selected the first 4PCs to include in the model by using the criteria that PCs had to have 0.15% more percent variation explained than the consecutive PC.

We chose to analyze the SNPs selected as cofactors by the maximum model as our top SNPs. The phenotypic variation explained by cofactor SNPs is calculated in the ‘mlmm’ package and corrects estimates for variation explained by fixed covariates. Manhattan plots were generated with the SNP p-values from the maximum model using ‘ggplot2’ (Wickham, 2016). The Bonferroni corrected significance threshold for p-values was calculated as 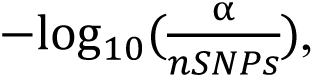 where α = 0.05 and *nSNPs* = 32703. Quantile-quantile plots in the supplemental figures were made with the ‘qqman’ package (Turner, 2018). SNPs selected as model cofactors by the maximum step model were further analyzed for their allelic effects and nearby genes. We used the Honeycrisp gene coordinates to overlap SNPs and genes as the genomes of this population were aligned to Honeycrisp v1.0 (Khan et al., 2022). Due to the low SNP coverage of our genome, we chose a gene linkage range of 25,000 base pairs as it is slightly larger than the 2,500 base pair global linkage average of *Malus* (Jung et al., 2020) but still relatively restricted. Candidate gene BLAST results and GO terms were obtained from GDR (Jung et al., 2019).

## 3. Results

Of the original 452 *Malus* accessions evaluated by Jurick II et al. (2011) for post-harvest rot in 2011, only 106 remain in the USDA-ARS collection. Low-pass sequencing performed by Khufu (Korani et al., 2021) mapped reads to the Honeycrisp v1.0 hap 1A genome with rates of 82.31% and 84.67% between the two multiplexed sample plates sent for sequencing. The Honeycrisp genome was chosen for its high contiguity and that many of the accessions included in this paper are hybrids between *M.* × *domestica* × *M. sieversii* or are classified as either of *M. domestica* two progenitor species, *M. sieversii* and *M. sylvestris* (Khan et al., 2022; Sun et al., 2020; Volk et al., 2022). Following the variant calling and imputation by Khufu, a total of 32,703 SNPs remained for GWAS. The average distance between SNPs was 38,270 base pairs.

After data curation, 96 *Malus* accessions were contained in the phenotype and genotype data. The *P. expansum* resistance categories of these accessions totaled 11 resistant, 22 moderately resistant, and 63 susceptible. The majority of the remaining 96 accessions were *M. domestica*, *M. sieversii*, and *M.* × *domestica* × *M. sieversii* hybrids (Figure 1). Aside from one accession of *M. sylvestris* and *M. prunifolia*, the moderate and completely resistant categories are composed of *M. sieversii* and hybrid accessions. Taxonomic classifications for each accession correspond to their current USDA Germplasm Resource Information Network (GRIN) classification as of October 2022 (USDA Agricultural Research Service, 2024; Volk et al., 2022).

**Figure 1.**
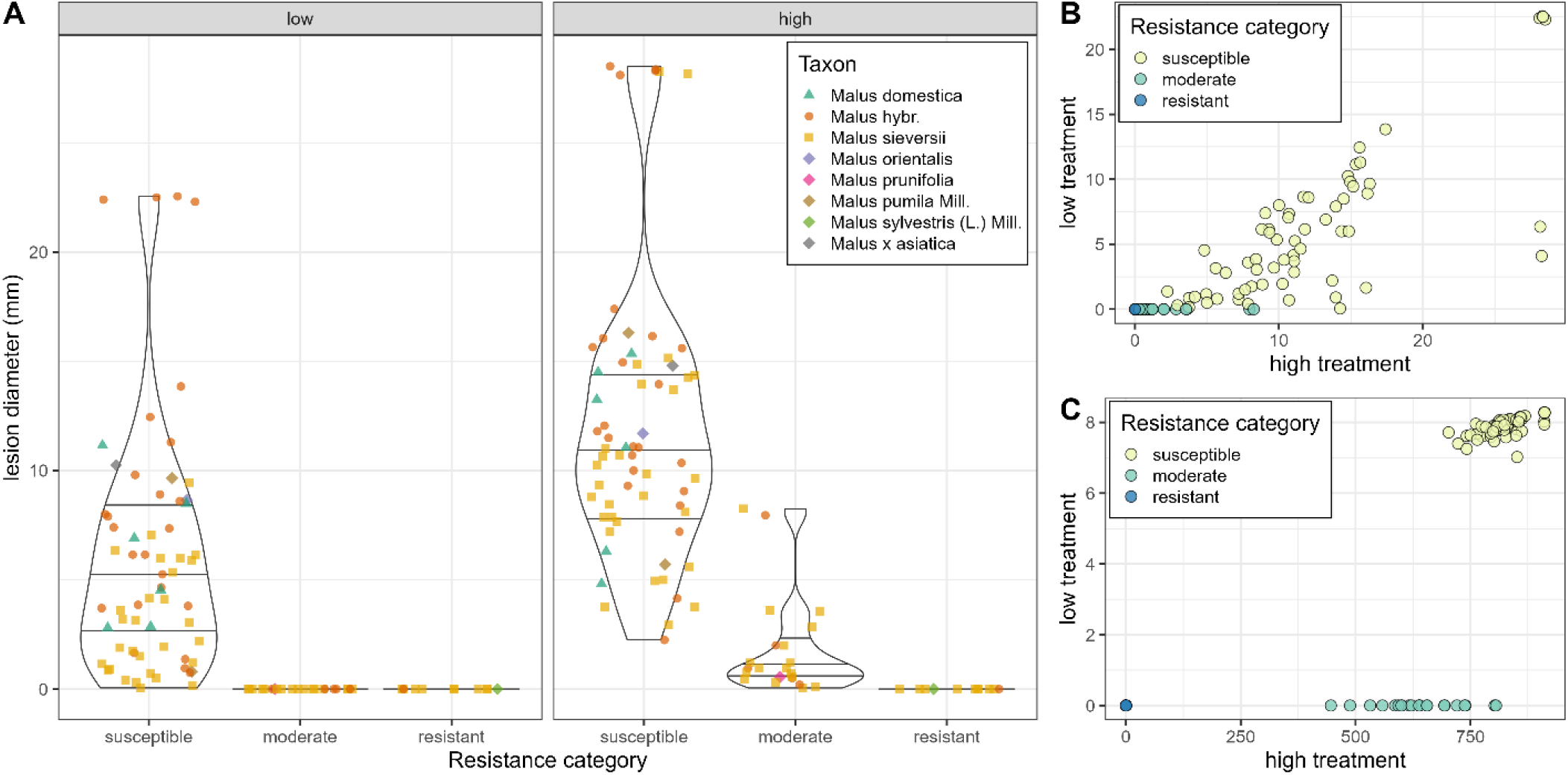
Summary of post-harvest rot of *Malus* accession when fruit is treated with low and high levels of *P. expansum* inoculum and categorized by their level of *P. expansum* resistance by Jurick II et al. (2011). **A)** Distribution of raw mean fruit lesion sizes of accessions colored by taxon, with different shapes distinguishing the main taxa: *Malus × domestica*, *Malus sieversii*, and hybrids, from the minor taxa in this population. **B)** Dimensions of the raw means, grouped by their *P. expansum* resistance category. **C)** Dimensions of normalized model-adjusted means taken as phenotypes for GWAS, grouped by their *P. expansum* resistance category.

Principal component analysis (PCA) and kinship conducted with the 32,703 SNPs in our genotype file showed little population structure (STable 5, SFigure 1). The first 4 PCs, which were included as covariates in our GWAS model, explained 12.53% of the genetic variance in this population (STable 5). PC1 alone is contributing half of that genetic variance (6.11%) and roughly divides the population by *Malus* taxa, especially the three main taxa which have significantly different PC1 values (Figure 2).

**Figure 2.**
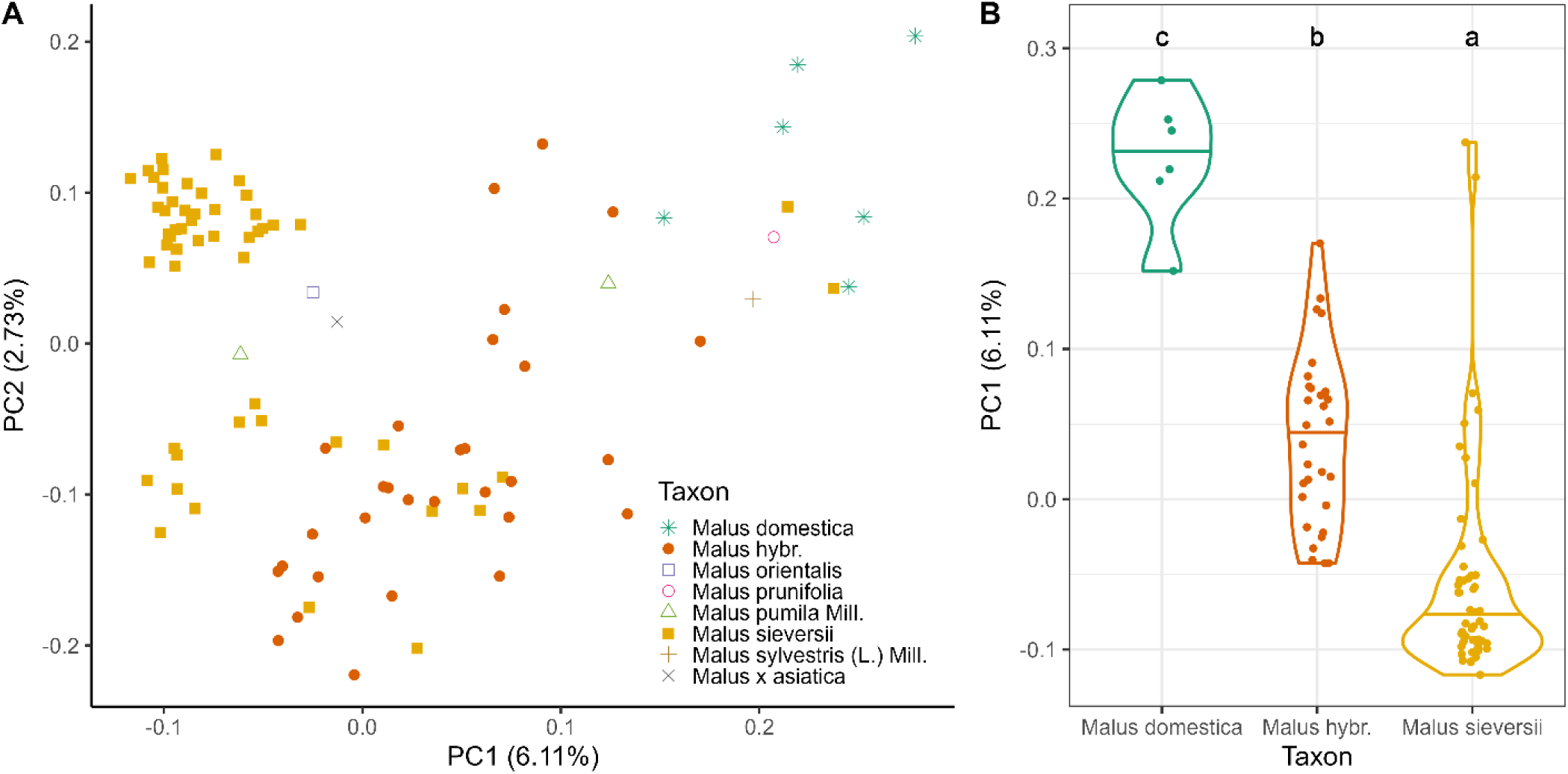
Principal component analysis of genotype data (32,703 SNPs). **A)** PC1 and PC2 colored by all Malus taxa used in the post-harvest rot GWAS. **B)** Distributions of PC1 values across the three main taxa in the population. Taxa not sharing any letter labeled at the top of the plot have PC1 values were determined significantly different by the Sidak-test (alpha 1 x 10^-4^).

### 3.1 Genome-wide association study

We used the multi-locus mixed model (MLMM) implemented by the ‘mlmm’ R package (Segura et al., 2012) to conduct GWAS for post-harvest rot lesion phenotypes under low (1 x 10^3^) and high (1 x 10^4^) *P. expansum* inoculum treatments individually and combined across treatments. For the high treatment phenotype, the maximum step, extBIC-optimized, and mBonf-optimized models selected three SNPs as cofactors at Chr05A_6378008, Chr07B_564388, and Chr17B_29483088 (Figure 3A, Table 1). These SNPs also passed the Bonferroni-corrected significance threshold (p < 1.53e^-06^), with the marker on Chromosome 7B exhibiting the strongest signal (*p* = 4.17e^-08^) and explaining 31.29% of the genetic variation. For the low treatment phenotype, the maximum model selected only one SNP at Chr03B_34046735 (Figure 1B) that explained 15.96% of the genetic variation. While the fact that neither the mbonf nor extBIC optimized model agreed on if this SNP was a cofactor, this location is within 2,908,876 base pairs of the *qM-Pe3.1* marker position when mapped to the Honeycrisp 1A genome (Luo et al., 2020). For the combined phenotype modeled across both treatments, the maximum and extBIC-optimized models selected the same SNPs as the high treatment phenotype. After GWAS, we did not explore the combined phenotype, since there was little variation in the explained variance and p-values of SNPs between the combined and high treatment phenotypes (STable 1-2).

**Figure 3.**
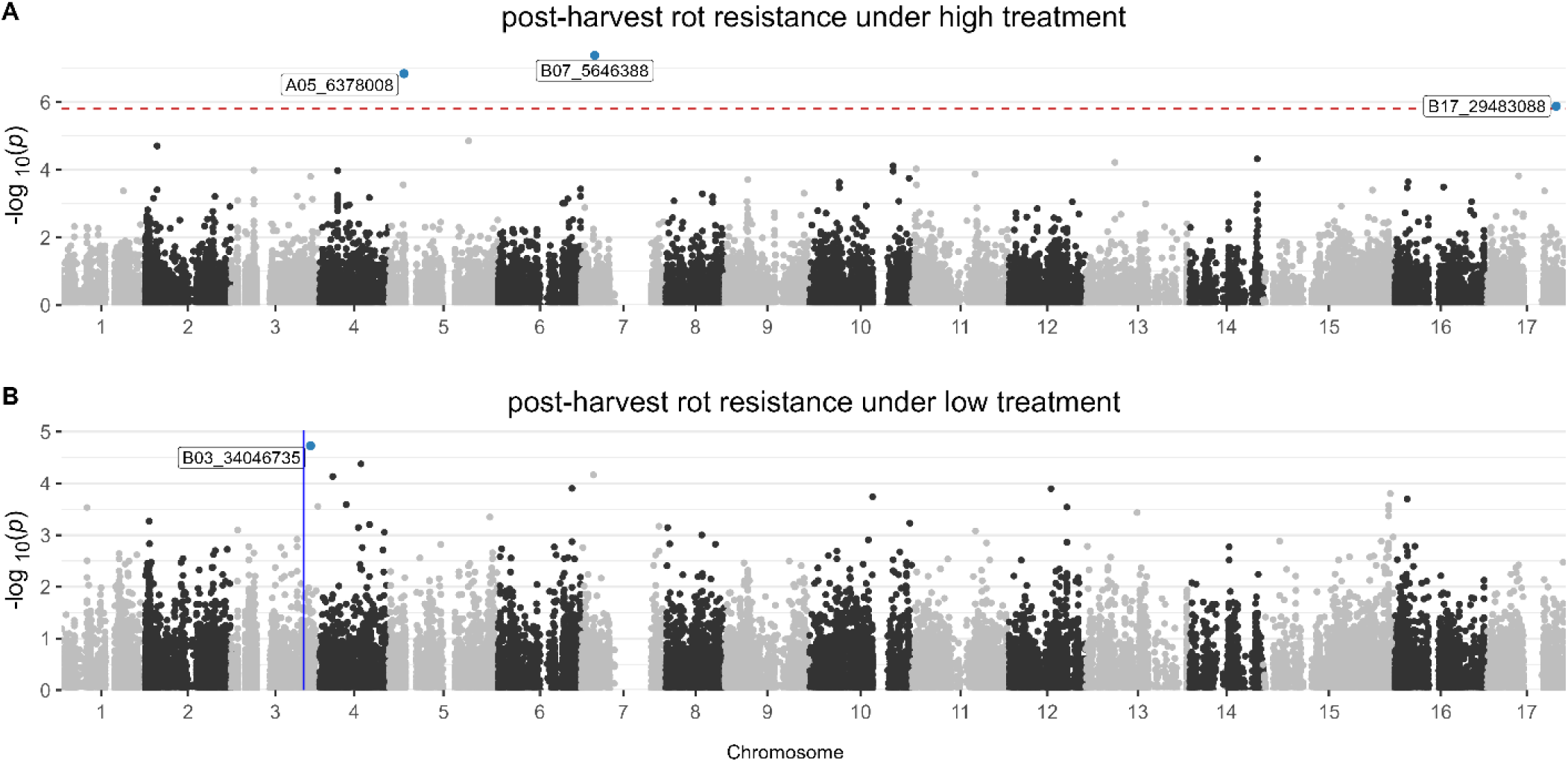
Manhattan plots of post-harvest rot resistance GWAS. Phenotypes are the diameter of rot lesions (mm) when fruits are treated with **(A)** high and **(B)** low levels of *P. expansum* inoculum. The red dotted line represents the Bonferroni corrected significance threshold (-log_10_*p >* 5.82). The blue horizontal line in panel B marks the *qM-Pe3.1* marker position.

**Table 1.**
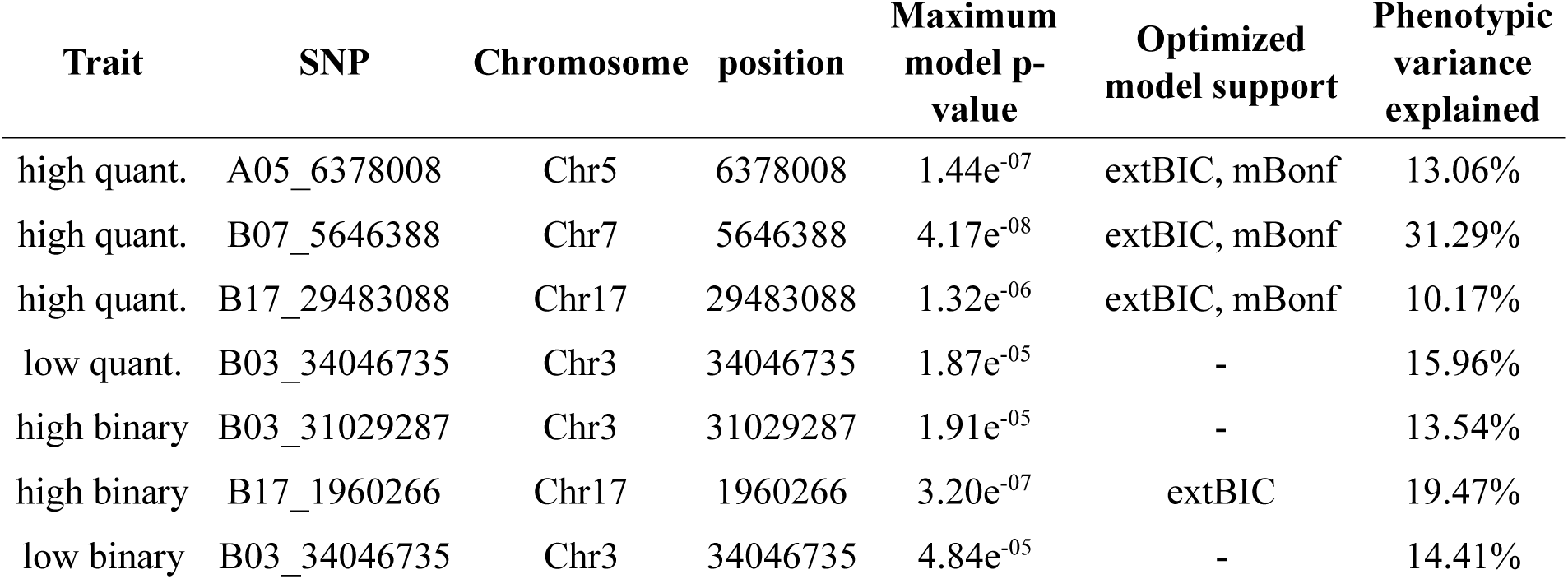
Top SNPs of MLMM GWAS with quantitative and binary phenotypes.

| Trait | SNP | Chromosome | position | Maximum model p-value | Optimized model support | Phenotypic variance explained |
| --- | --- | --- | --- | --- | --- | --- |
| high quant. | A05_6378008 | Chr5 | 6378008 | 1.44e<sup>-07</sup> | extBIC, mBonf | 13.06% |
| high quant. | B07_5646388 | Chr7 | 5646388 | 4.17e<sup>-08</sup> | extBIC, mBonf | 31.29% |
| high quant. | B17_29483088 | Chr17 | 29483088 | 1.32e<sup>-06</sup> | extBIC, mBonf | 10.17% |
| low quant. | B03_34046735 | Chr3 | 34046735 | 1.87e<sup>-05</sup> | - | 15.96% |
| high binary | B03_31029287 | Chr3 | 31029287 | 1.91e<sup>-05</sup> | - | 13.54% |
| high binary | B17_1960266 | Chr17 | 1960266 | 3.20e<sup>-07</sup> | extBIC | 19.47% |
| low binary | B03_34046735 | Chr3 | 34046735 | 4.84e<sup>-05</sup> | - | 14.41% |

Additionally, we conducted GWAS with the binary phenotype data to compare the difference in sensitivity between qualitative and quantitative datasets (STable 1-2). In the high treatment phenotype, the maximum model selected two SNPs as cofactors: one at Chr17B_1960266 that was also selected by the extBIC-optimized model, and one at Chr03B_31029287 (SFigure 2A). While the Chr03 SNP was not chosen by the extBIC model and did not pass the Bonferonni threshold (*p* = 1.91e^-05^), it is 108,572 base pairs away from the *qM-Pe3.1* marker located at Chr03_31137859 (Luo et al., 2020). In the low treatment phenotype, the same SNP selected by the quantitative low phenotype at Chr03B_34046735 was selected by the model (SFigure 2B). However, the binary data assigned it a lower *p*-value and portion of phenotypic variance explained than the phenotype in the quantitative dataset (Table 1, STable 2).

### 3.2 SNP allele effects

Most model-selected SNPs show significant differences in lesion size between the two homozygous allele pairs, and a few have many resistant *Malus* accessions under the allele with significant reduction in lesion diameter (Figure 4, SFigure 4). We also saw some alleles respond differently depending on the inoculum concentration. At SNP B03_31029287 from the binary phenotype under the high inoculum treatment, the “G” allele homozygous accessions have reduced lesion size compared to the heterozygotes and “A” allele homozygotes under low inoculum concentrations. Yet under high *P. expansum* concentrations, accessions with one or two “G” alleles have the same lowered phenotype. In the low treatment SNP B03_34046735, we observe the “CC” genotype contains all the resistant accessions and the majority of moderately resistant accessions and is significantly lower under the high inoculum treatment (SFigure 4). B03_34046735 also has only one accession with the “AA” genotype, and this accession was categorized as susceptible by Jurick II et al. (2011). It is possible that the replicates from this accession may not fully reflect the phenotype of all “AA” accessions in the whole population. SNPs A05_6378008 and B07_5646388 both have “A” alleles significantly linked to lower lesion diameters under both treatments, with the “AA” genotype having the strongest phenotype (Figure 4C&D, SFigure 4). They also appear to be the SNPs of rarer alleles, as few accessions have the “AA” call at these SNPs (SFigure 4). At SNP B17_1960266 from the binary data’s high treatment phenotype, the “AA” allele contains all but one of the resistant accessions and has significantly lower phenotypes than the “GG” and heterozygous genotypes (SFigure 4). Lastly, SNP B17_29483088 from the high treatment phenotype in the quantitative data shows no difference between alleles under the low inoculum treatment. However, when treated with high concentrations of *P. expansum* inoculum, there is a significant reduction in lesion size for accessions with the “AA” genotype (Figure 4F). While the heterozygous phenotype is closer to the “AA” phenotype than the “GG” phenotype, it was not significantly different than either homozygous genotype as determined by the Sidak-test (SFigure 4).

**Figure 4.**
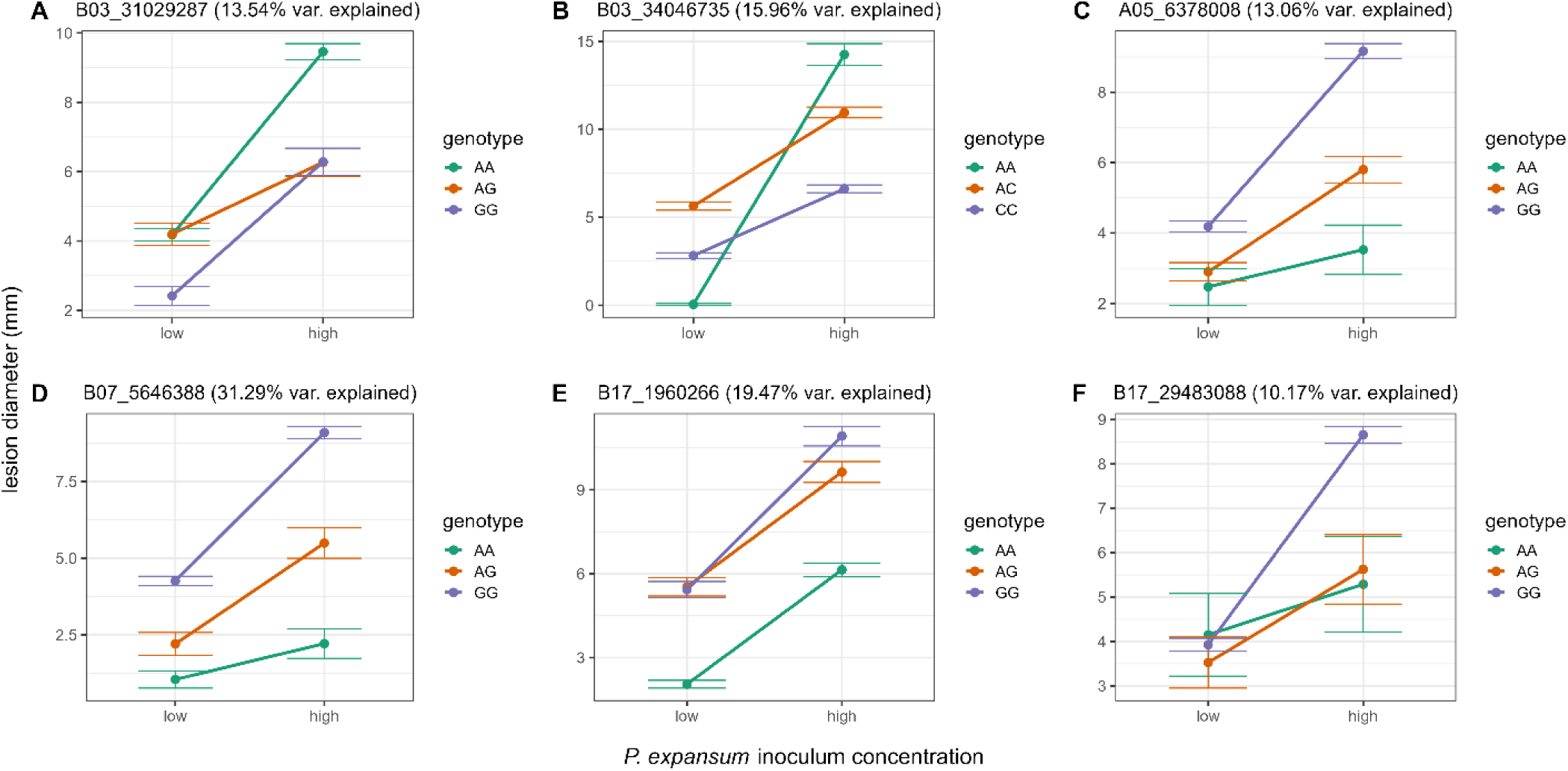
Allelic effects of model-selected SNPs under low and high concentrations of *P. expansum* inoculum treatment. Even if SNPs were detected by models with binary phenotypes, plots are shown with quantitative phenotypes to improve visualization. Points represent average phenotype of each allele pair and error bars represent standard error. SNPs are from GWAS modeled with the **A,E)** high binary phenotype and **B)** low quantitative phenotype (SNP B03_34046735 is also a cofactor in the low binary phenotype model and explained 14.4% of phenotypic variance), and **C,D,F)** high quantitative phenotype.

### 3.3 Candidate genes

Using a linkage range of ±25kb and comparing SNPs to genes on either haplotype, we found 73 genes linked to the six SNPs selected by the MLMM GWAS models (STable 3 and 4). We have highlighted candidate genes from the quantitative phenotypes (Table 2) with annotations related to our current understanding of the post-harvest rot phenotypes. For the low treatment phenotype on chromosome 3 we found 4 genes annotated as flavin-containing monooxygenase family proteins, an ortholog to DEGP protease 2 in *Arabidopsis thaliana*, and a cell wall/vacuole inhibitor of fructosidase 1. For the high treatment phenotype, we found a NAC containing protein, a non-intrinsic ABC protein and an uncharacterized gene with a gene ontology (GO) term for innate immune response linked to the SNP on chromosome 5. The uncharacterized gene status is determined by the GDR BLAST results of all genes within its databases from different sources (Jung et al., 2019). However, we performed a protein-protein BLAST of this sequence to the non-redundant protein database (Camacho et al., 2009) and found additional annotations. Most top hits were uncharacterized, but the few characterized proteins were a mucus-7-like protein in valley oak (XM_031100062), pep-1 precursor in *Prunus domestica* (UNA27434), and another pep precursor (PROPEP4) in pear (DAD54771; Ruiz et al. 2018). On chromosome 7, the SNP for high treatment phenotypes is located inside a gene’s intron on each haplotype: Maldo.hc.v1a1.ch7A.g40987 is annotated as a magnesium ion binding protein (STable 3) and Maldo.hc.v1a1.ch7B.g43072 is annotated as a RAC-like protein. Other chromosome 7 notable hits are the glutaredoxin family protein and COBRA-Like 7 gene. Lastly, on chromosome 17 the SNP for high treatment phenotypes is linked to a locus receptor-like (LRL) protein kinase-like gene associated with leaf rust resistance and a pathogenesis-related 5-like (PR5) receptor kinase.

**Table 2.** Functional annotation of candidate genes linked to top SNPs in GWAS with quantitative phenotypes. BLAST results and GO terms obtained from Genome Database for Rosaceae.

| Trait | SNP | p-value | Gene | BLAST results | GO terms |
| --- | --- | --- | --- | --- | --- |
| low | B03_3404673<br>5 | 1.87E<sup>-05</sup> | Maldo.hc.v1a1.ch3B.g33434 | Araport11: AT1G47960.2 \| cell wall / vacuolar inhibitor of fructosidase 1 | enzyme inhibitor activity (GO:0004857)<br>negative regulation of catalytic activity (GO:0043086) |
| low | B03_3404673<br>5 | 1.87E<sup>-05</sup> | Maldo.hc.v1a1.ch3A.g31820 | Araport11: AT1G48910.1 \| Flavin-containing monooxygenase family protein<br>NCBI_nr: probable indole-3-pyruvate monooxygenase YUCCA10 [Malus domestica] | flavin adenine dinucleotide binding (GO:0050660)<br>NADP binding (GO:0050661) |
| low | B03_3404673<br>5 | 1.87E<sup>-05</sup> | Maldo.hc.v1a1.ch3A.g31819 | Araport11: AT1G48910.1 \| Flavin-containing monooxygenase family protein | flavin adenine dinucleotide binding (GO:0050660)<br>NADP binding (GO:0050661) |
| low | B03_3404673<br>5 | 1.87E<sup>-05</sup> | Maldo.hc.v1a1.ch3A.g31818 | Araport11: AT1G48910.1 \| Flavin-containing monooxygenase family protein | flavin adenine dinucleotide binding (GO:0050660)<br>NADP binding (GO:0050661) |
| low | B03_3404673<br>5 | 1.87E<sup>-05</sup> | Maldo.hc.v1a1.ch3A.g31817 | Araport11: AT2G47940.1 \| DEGP protease 2<br>NCBI_nr: protease Do-like 2, chloroplastic [Malus domestica] | protein binding (GO:0005515)<br>proteolysis (GO:0006508)<br>serine-type endopeptidase activity (GO:0004252) |
| high | A05_6378008 | 1.44E<sup>-07</sup> | Maldo.hc.v1a1.ch5B.g36771 | Araport11: AT1G63270.1 \| non-intrinsic ABC protein 10 | ATP binding (GO:0005524)<br>cytochrome complex assembly (GO:0017004)<br>transmembrane transporter activity<br>(GO:0022857) |
| high | A05_6378008 | 1.44E<sup>-07</sup> | Maldo.hc.v1a1.ch5B.g36770 | Uncharacterized protein | innate immune response<br>(GO:0045087) |
| high | A05_6378008 | 1.44E<sup>-07</sup> | Maldo.hc.v1a1.ch5A.g34880 | Araport11: AT1G34190.1 \| NAC domain containing protein 17<br>NCBI_nr: NAC domain-containing protein 17-like [Malus domestica] | DNA binding<br>(GO:0003677)<br>regulation of DNA-templated transcription<br>(GO:0006355) |
| high | B07_5646388 | 4.17E<sup>-08</sup> | Maldo.hc.v1a1.ch7B.g43072 | Araport11: AT5G62880.2 \| RAC-like 10 | GTPase activity<br>(GO:0003924)<br>GTP binding<br>(GO:0005525)<br>small GTPase-mediated signal transduction<br>(GO:0007264) |
| high | B07_5646388 | 4.17E<sup>-08</sup> | Maldo.hc.v1a1.ch7B.g43069 | Araport11: AT4G12560.3 \| F-box and associated interaction domains-containing protein | protein binding<br>(GO:0005515) |
| high | B07_5646388 | 4.17E<sup>-08</sup> | Maldo.hc.v1a1.ch7B.g43074 | NCBI_nr: putative COBL7 (COBRA-LIKE 7) [Malus domestica] |  |
| high | B07_5646388 | 4.17E<sup>-08</sup> | Maldo.hc.v1a1.ch7A.g40986 | Araport11: AT5G01420.1 \| Glutaredoxin family protein<br>NCBI_nr: uncharacterized protein At3g28850-like isoform X1 [Malus domestica] |  |
| high | B07_5646388 | 4.17E<sup>-08</sup> | Maldo.hc.v1a1.ch7A.g40985 | Araport11: AT5G20230.1 \| blue-copper-binding protein<br>NCBI_nr: cucumber peeling cupredoxin-like [Malus domestica] | electron transfer activity<br>(GO:0009055) |
| high | B07_5646388 | 4.17E<sup>-08</sup> | Maldo.hc.v1a1.ch7A.g40984 | Araport11: AT5G22060.1 \| DNAJ homologue 2<br>NCBI_nr: dnaJ protein homolog [Malus domestica] | ATP binding<br>(GO:0005524)<br>heat shock protein binding<br>(GO:0031072)<br>Hsp70 protein binding<br>(GO:0030544)<br>protein folding<br>(GO:0006457)<br>response to heat<br>(GO:0009408)<br>unfolded protein binding<br>(GO:0051082) |
| high | B17_2948308<sub>8</sub> | 1.32E<sup>-06</sup> | Maldo.hc.v1a1.ch17A.g23854 | Araport11: AT5G38260.1 \| Protein kinase superfamily protein<br>SwissProt: LRL21_ARATH LEAF RUST 10 DISEASE-RESISTANCE LOCUS RECEPTOR-LIKE PROTEIN KINASE-like | polysaccharide binding<br>(GO:0030247) |
| high | B17_2948308<sub>8</sub> | 1.32E<sup>-06</sup> | Maldo.hc.v1a1.ch17B.g24983 | NCBI_nr: 2-hydroxyacyl-CoA lyase [Malus domestica]<br>Araport11: AT5G17380.1 \| Thiamine pyrophosphate dependent pyruvate decarboxylase family protein | catalytic activity<br>(GO:0003824)<br>magnesium ion binding<br>(GO:0000287)<br>thiamine pyrophosphate binding<br>(GO:0030976) |
| high | B17_2948308<br>8 | 1.32E-06 | Maldo.hc.v1a1.ch17A.g23855 | Araport11: AT5G38280.1 \| PR5-like<br>receptor kinase<br>NCBI_nr: PR5-like receptor kinase<br>isoform X3 [Malus domestica] | ATP binding<br>(GO:0005524)<br>protein kinase activity<br>(GO:0004672)<br>protein phosphorylation<br>(GO:0006468) |
| high | B17_2948308<br>8 | 1.32E-06 | Maldo.hc.v1a1.ch17B.g24990 | Araport11: AT1G66730.1 \| DNA<br>LIGASE 6 | ATP binding<br>(GO:0005524)<br>DNA binding<br>(GO:0003677)<br>DNA biosynthetic<br>process (GO:0071897)<br>DNA ligase (ATP)<br>activity (GO:0003910)<br>DNA recombination<br>(GO:0006310)<br>DNA repair<br>(GO:0006281) |

We also wanted to highlight some interesting findings from the GWAS model for high treatment phenotypes in the binary data (Table 3; STable 4). Most notably are the putative disease resistance protein (RAG3) gene, a gene encoding a LOB domain-containing protein, and an open reading frame DNA fragment spanning a resistance locus for *Erwinia amylovora* (Fahrentrapp et al., 2013). We also found genes with GO terms that suggest minor roles in pathogen defense, such as lyase, ATP hydrolysis, and glycosyltransferase activity.

**Table 3.**
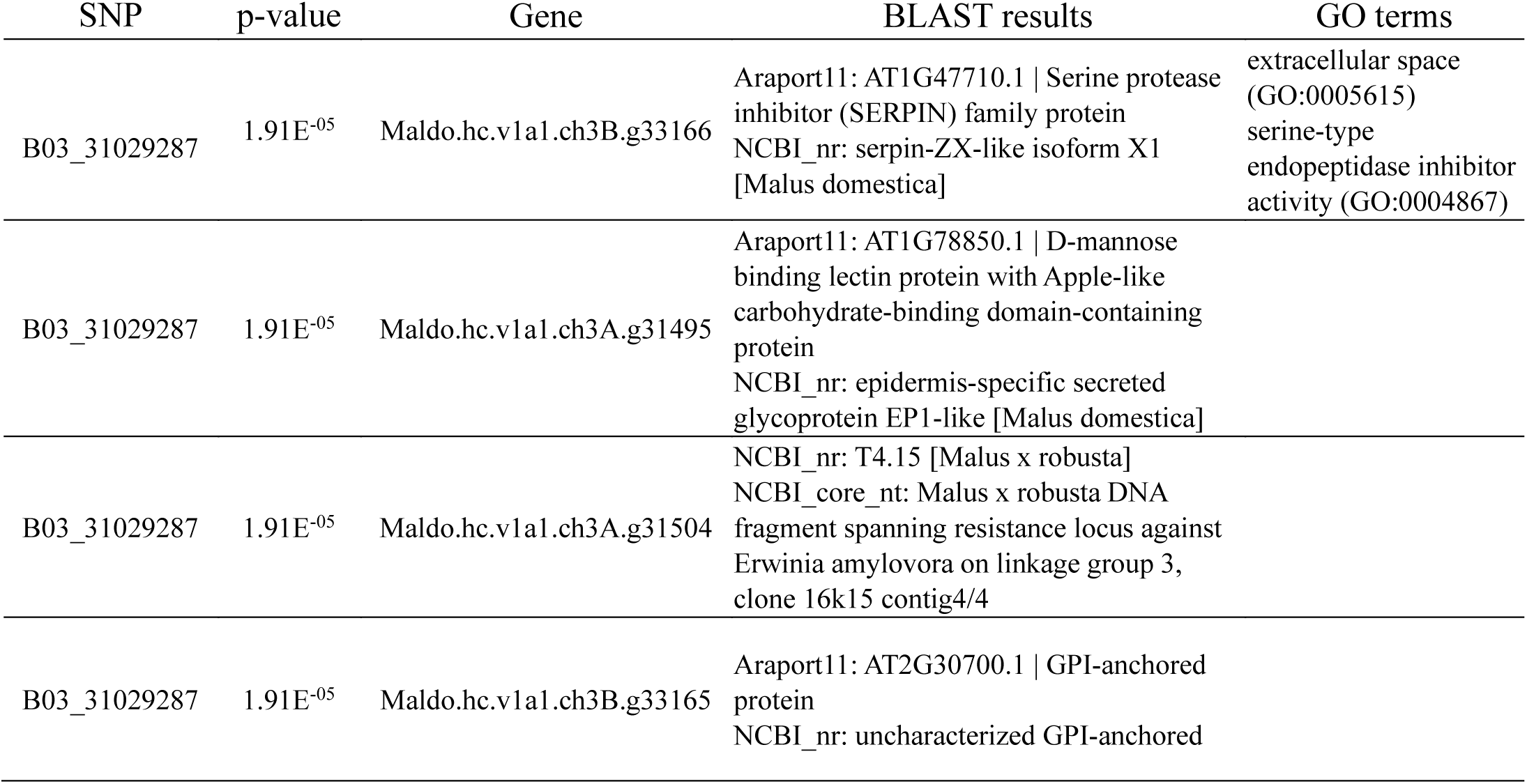
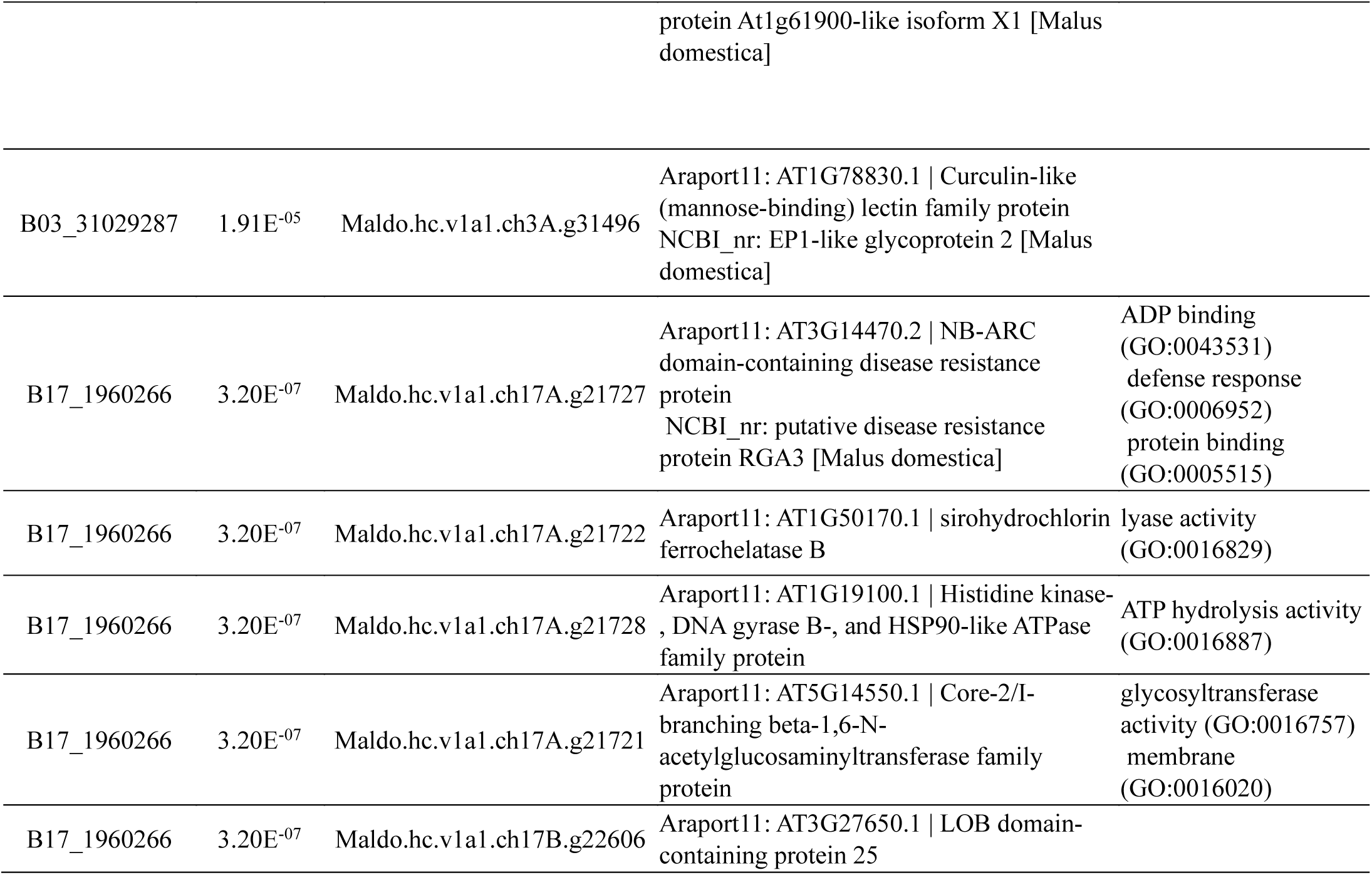
Functional annotation of candidate genes linked to SNPs in the high treatment with binary phenotypes.

## 4. Discussion

We have demonstrated that the application of new technologies and methods to existent, historical datasets can yield new insights and targets for breeding programs. Despite challenges in recovering DNA samples from all original *Malus* accessions and using a low-pass sequencing approach, we were able to generate a sufficient and meaningful genetic dataset suitable for association mapping with the phenotypes previously reported by Jurick II et al. (2011). The kinship analysis (SFigure 1) and SNP PCA of our 96 accessions and 32,703 SNPs signaled that this dataset was diverse enough for a GWAS. To our knowledge, this is the first work using low-pass sequencing coupled with GWAS in *Malus spp.* However, this approach has been successfully implemented in other crops such as dry bean, millet, mustard, and spinach to genotype populations and identify variants associated with disease resistance (Bhattarai et al., 2023; Khound et al., 2022; Rana et al., 2019; Simons et al., 2021).

### 4.1 Low-pass sequencing results in potential false negatives and inability to determine exact causal genes

The SNP density is low compared to the estimated 2,500 base pair linkage disequilibrium decay in *Malus domestica* (Jung et al., 2020). Thus, the SNPs significantly associated with phenotypic variation (Table 1) are likely to be approximate, rather than exact, indicators of causal genes. It is also very possible this weakened the signal of many true hits, as we observe additional peaks on chromosomes 2, 5, and 14 in the high treatment GWAS approaching the Bonferroni threshold (Figure 3A). We widened our window for linked candidate genes to 25,000 kb to account for the low SNP coverage and false negatives while remaining below the average SNP distance of 38,270 bp to retain a relatively narrow scope. Within these windows, we found many promising candidate genes for post-harvest rot resistance (Table 2), though, like in many GWAS, the exact causal gene cannot be confirmed. Moreover, the observation of numerous significant SNPs across four chromosomes suggests that high resistance/immunity to *P. expansum* is a quantitative trait. Nonetheless, these loci provide promising targets to further fine-map for the causal genes involved with *P. expansum* post-harvest rot resistance.

### 4.2 Treatments using high pathogen inoculum concentrations increased GWAS signal

The phenotypes under low concentrations of *P. expansum* inoculum had low signal in GWAS, with only one SNP, B03_34046735, detected regardless of the type of phenotype data used. Additionally, only the SNPs found in the high treatment GWAS were supported by both optimized models (STable 2). We hypothesized that there would be many SNPs linked to rot resistance in the low treatment GWAS, as there are likely many loci conferring broad spectrum pathogen resistance. Complete resistance, as observed in the few *M. sieversii, M. sylvestris* and hybrid accessions, was hypothesized to be under fewer, strong-effect loci containing resistance genes specific to *P. expansum*. Especially with the existence of the moderately resistant accessions, there should be mechanisms conferring resistance in the low treatment, even if it is broken under the high treatment (Figure 1). One possible explanation is the lack of phenotypic representation of the population since only 96 lines of the original study were recovered. There is clear phenotypic variation in this *Malus* population (Figure 1), and if there are many genetic factors contributing sufficient resistance when stressed with low inoculum concentrations, we may not be capturing enough accessions with alleles for each type of low-level resistance. For example, we can see a SNP (B07_5108632) with moderate *p*-value in the same region as the B07_5646388 SNP in the high treatment data (Figure 3), and with more accessions we might see the signal improve. Unfortunately, many of the original accessions have been removed from the USDA *Malus* collection since these phenotypes were collected.

As expected, most SNPs had alleles with overall better resistance under both treatments. For some of the SNPs, the optimal allele for rot resistance is only detected under the high *P. expansum* inoculum treatment as there is little difference between allele effects in the low treatment (Figure 4C,F). SNP B03_31029287 has an optimal genotype “GG” that is more resistant than other genotypes under both treatments (Figure 4A). Interestingly, the heterozygote accessions have a dosage or threshold effect where they are more like the “AA” allele under the low treatment, but under the high treatment they have the same phenotype as “GG” accessions. SNP B03_31029287 is also located within ∼100Kb of the *qM-Pe3.1* blue mold resistance marker from Luo et al. (2020). Our SNP calls have PI 613981 with an imputed “GG” call, which is associated with the resistant phenotype (STable 6). Taken together, our results support the use of *qM-Pe3.1* derived from PI 613981 as a marker for resistance. However, our dataset was not limited by the narrow genetics of a mapping population such as the PI 613981 × ‘Gala’ used by Norelli et al. (2017).

### 4.3 Quantitative measurements are highly informative and binary data is useful

The phenotypic data from Jurick II et al. (2011) provided the opportunity to test two different methods of phenotypic measurement. With a generalized linear model and binomial distribution, we were able to generate estimated means from the 20 fruit replicates in both treatments for GWAS and detected three significant SNPs across both treatments. We are confident in the usefulness of the binary phenotypes, as they identified SNPs within the same genetic regions as the quantitative phenotypes (Figure 3). In the low treatment, the binary and quantitative phenotype GWAS even selected the same SNP (B03_34046735) as a model cofactor and attributed similar explained phenotypic variance to the SNP (Table 1). Binary presence/absence categorization of post-harvest rot in apples has been standard since the 1900s (Horsfall and Barratt, 1945). We wanted to include GWAS results with the binary phenotypes to show that other historic datasets of curated plant collections may also be candidates for re-analysis with association or joint-linkage mapping. These methods are also applicable to other phenotypes in horticultural research that are quantitative in nature but are limited to qualitative classifications due to the complex nature of obtaining quantitative measurements. One example is phenotyping fruit bruising sensitivity during shipment, which is a complex trait influenced by fruit morphology and multiple biochemical processes (K. Mitsuhashi-Gonzalez et al., 2010) with phenotypic variance across apple accessions (Ericsson and Tahir, 1996).

When possible, quantitative measurements are logically preferred over binary, as they contain more information about phenotypic variation. In GWAS, we have shown that the quantitative phenotypes identify more SNPs with stronger phenotypic associations compared to their binary counterparts (Table 1). However, we would like to highlight that while the quantitative data is an improvement over the binary categorizations of disease, the measurements in the historic data were taken as whole millimeters or “count data” (Jurick II et al., 2011). Unless the scale of measurements is large enough to act as linear data, count measurements are best modeled with different distributions, like the negative binomial distribution we used, than those used in standard linear regressions (Zeileis et al., 2008). Traits like days to harvest or flower counts are examples of typical count phenotypes, but post-harvest rot lesion size is a continuous phenotype. If we were able to obtain precise lesion measurements, we hypothesize that these phenotypes would improve the signal to detect additional significant SNPs like the quantitative phenotypes improved over the binary. Advancements like high-throughput image-based phenotyping (Gehan et al., 2017; Rueden et al., 2017) can make this transition feasible for traits like post-harvest rot that are traditionally time and labor intensive.

### 4.4 GWAS identifies disease resistance loci

Within a 25kb area of the top SNPs associated with the resistant wild apple accessions, many candidate plant defense genes were discovered (Table 2-3). While their direct role in resistance to blue mold decay has not been demonstrated, these genes provide an exciting opportunity to further explore their role in decay resistance (Jurick II, 2022). Some candidates in the apple genome have known annotations and corresponding functional work describing their role in disease resistance in model and other crop plants that could be extrapolated to our system. To categorize our results, we observed three main classes of proteins encoded by the candidate genes that comprise A) enzymes (*e.g.,* flavin monooxygenase) B) signaling components (*e.g.,* GPI-anchored protein) and C) hallmark plant resistance genes (*e.g.,* PR5 and LRR). We have chosen not to discuss all the identified loci here in the discussion but have focused on ones of primary interest for their well characterized roles in plant defense.

Two genes (e.g. Maldo.hc.v1a1.ch7B.g43074 [Cobra-like 7], Maldo.hc.v1a1.ch3A.g31818 [Flavin-containing monooxygenase] identified in this study have interesting parallels with previous findings in the apple-blue mold system and with other fungal pathosystems. A recent study conducted by Moriya et al. (2019) found that Cobra-like genes were strongly associated with resistance to Alternaria blotch in apple, caused by the necrotrophic fungal pathogen *Alternaria alternata*. COBRA-Like (COBL) genes encode glycosylphosphatidylinositol (GPI) anchored polypeptides that are key regulators in the orientation of cell expansion and cellulose crystallinity (Niu et al., 2015). Interestingly, studies with the same wild apple germplasm accessions have shown that wound healing is a primary component of blue mold resistance (Janisiewicz et al., 2016). Hence, it is plausible that this locus and other cell-wall associated biosynthetic genes are involved in host defense responses in *Penicillium*-wild apple fruit interactions. Additionally, a gene encoding a flavin-containing monooxygenase, associated with phenolic metabolism and plant defense, was found. In tomato and Arabidopsis, the *FMO1* locus that encodes a flavin monooxygenase that modulates ROS homeostasis and regulates phenolic compound production correlated with antimicrobial compounds (Koch et al., 2006; Wang et al., 2023). Curiously, we observed a link between the presence of the Maldo.hc.v1a1.ch3A.g31818 locus that encodes a flavin monooxygenase in wild apple fruit with our previous data showing that the wild apple accessions resistant to blue mold have increased levels of phenolics and high ROS accumulation (Janisiewicz et al., 2016; Janisiewicz and Jurick, 2017). Hence, specific genes, like Maldo.hc.v1a1.ch3A.g31818, could be responsible for the production of antimicrobial phenolic compounds and ROS in wild apples with resistance to blue mold. However, definitive evidence concerning their roles would need to be conducted using a combination of functional genomic, metabolomic and biochemical approaches.

Genes encoding well characterized disease resistance loci were found to encode a LRL receptor (LEAF RUST 10 DISEASE-RESISTANCE LOCUS RECEPTOR-LIKE PROTEIN KINASE-like) and one a PR5-like receptor kinase. Genes modulating disease resistance have been discovered and studied in model and economically important plant species (Ding et al., 2022). The LRL encodes a serine/threonine receptor-like kinase that are involved in a wide variety of plant functions including: anther and ovule development, pollen-pistil interactions, vascular tissue development, as well as in leaf rust resistance and pathogen triggered immunity (Jose et al., 2020). The presence of this LRL is of major biological and translative potential as this gene could be edited in apple to make fruit more resistant to blue mold decay. Additionally, on the pathogen side, the cognate *P. expansum* elicitor that interacts with the apple LRL can also be determined via functional genetic and omics-based modalities that may lead to effective decay control strategies. Pathogenesis-related protein (PR) 5 is a thaumatin-like protein that is central to the systemic acquired resistance (SAR) response. PR5 is expressed along with other PR proteins, such as PR1 and PR2, during SAR and the encoded protein itself has antifungal activity (El-kereamy et al., 2011). Interestingly, expression of PR5 homologues from *Prunus domestica* was found to correlate with resistance to brown rot fungi. el-Kereamy et al. (2011) showed that the *PdPR5-1* gene has a role in activating other defense pathways, including phytoalexin production in plum. Therefore, PR5 overexpression in apple would be an exciting target to test for enhanced resistance to the necrotrophic blue mold fungus *P. expansum*. The putative resistance gene homologs RGA3 (Maldo.hc.v1a1.ch17A.g21727) and Maldo.hc.v1a1.ch3A.g31504 have roles in pathogen recognition and response in other organisms. RGA3 has been found to confer broad spectrum resistance to potato late blight caused by *Phytophthoria infestans*, a hemibiotrophic fungus with a necrotrophic phase in later stages of the disease (Song et al., 2003). Maldo.hc.v1a1.ch3A.g31504 BLASTs to an open reading frame homologous to a RGA in a study of fire-blight resistance candidate genes in apples and pears (Fahrentrapp et al., 2013). Fahrentrapp et al. (2013) identified their candidate (T4.15) as most similar to a TIR NBS-LRR RGA in peanut (*Arachis hypogaea*). While fire blight is caused by the bacteria *Erwinia amylovora*, it’s possible that this gene also aids broad spectrum recognition and resistance, like RGA3.

While our study shows new genes associated with resistance to blue mold decay in wild apple fruit, our findings should be considered significant, but cursory given the fact that we have no functional data at present. However, the well annotated loci in apple that we discovered using GWAS remain impactful both biologically and translationally. We envision that a functional approach via CRISPR and or traditional breeding to move these loci into commercial apple cultivars will be of great value to reduce blue mold decay in the future. Since all commercial apple cultivars are susceptible to blue mold, improved cultivars with decay resistance would be a welcomed tool by industry to aid in decay management. Our findings highlight the importance of studying germplasm to identify new sources of genetic resistance and understand host resistance mechanisms, and demonstrate the power of GWAS in deducing genes associated with blue mold resistance that would not be possible using only susceptible commercial apple cultivars (Jurick II et al., 2011).

### 4.5 Incorporating findings into breeding programs

In our study, we identified an additional novel four loci (excluding the two near *qM-Pe3.1*). These additional loci can provide potentially novel resistance from different sources than PI 613981 derived resistance (STable 6). For example, PI 687735 and PI 613981 carry non-imputed desirable alleles for the SNPs B17_1960266 and A05_6378008. These two SNP’s additionally explain 10.17% and 13.06% of the variation, a similar amount as the Chr03 loci identified in our study and in Norelli et al. (2017). The significant SNP B07_5646388 explains the greatest percent of variation (>30%). Of the resistant lines genotyped in our study, PI 657038 and PI 687735 carry non-imputed desirable alleles for this SNP (STable 6). These accessions provide opportunities for pyramiding resistance associated loci, resulting in more durable blue mold resistance. Since our results indicate polygenic resistance, taking a multi-locus, marker-assisted selection or genome selection approach could accelerate the introgression of these alleles. These methods have already been utilized in apple breeding for blue mold resistance and fruit quality traits (Kostick et al., 2023; Kumar et al., 2012; Luo et al., 2020; Muranty et al., 2015). Ultimately, breeding for blue mold resistance will increase the sustainability and profitability of apple production. Moreover, host-resistance can open new marketing opportunities for apple growers by reducing the usage of fungicides that are restrictive for export markets.

## Supporting information

STable 1

STable 2

STable 3

STable 4

STable 5

STable 6

## 5. Data Availability

The variant called haplomap and imputed haplomap have been deposited into the Genome Database for Rosaceae (Jung et al., 2019). The *Malus* germplasm accessions used in this study can be requested through the USDA-ARS National Plant Germplasm System (NPGS) GRIN-Global website (USDA Agricultural Research Service, 2024).

## 6. Author contributions: CRediT

LW contributed to the development of methodology, conducted the formal analysis, interpreted the results, writing of the original draft, and conducted the reviewing and editing. Co-authors JSB, TC, BE, DR, BG, WJ all contributed to resource acquisition, conducting experiments, and reviewing and editing the manuscript. Corresponding authors WMJ II and CG conceptualized and led the investigation, developed the methodology, provided supervision, interpreted the results, writing of the original draft, and conducted the reviewing and editing.

## 8. Supplemental Figures

**STable 1. Model phenotypes used in GWAS and metadata from inoculation experiment.**

**STable 2. Extended SNP (p < 1e-4) and RSS output from MLMM models.**

**STable 3. Full candidate gene list from quantitative measures.** Distance is the base pairs between SNPs and genes.

**STable 4. Full candidate gene list from binary measures. Distance is the base pairs between SNPs and genes.**

**STable 5. Genotypic variance explained by PCs 1-4.**

**STable 6. Resistant accessions and their genotype call for each significant SNP/trait association.**

Bold alleles signify the desirable allele and was non-imputed. Italics signify imputed SNP calls.

**SFigure 1.**
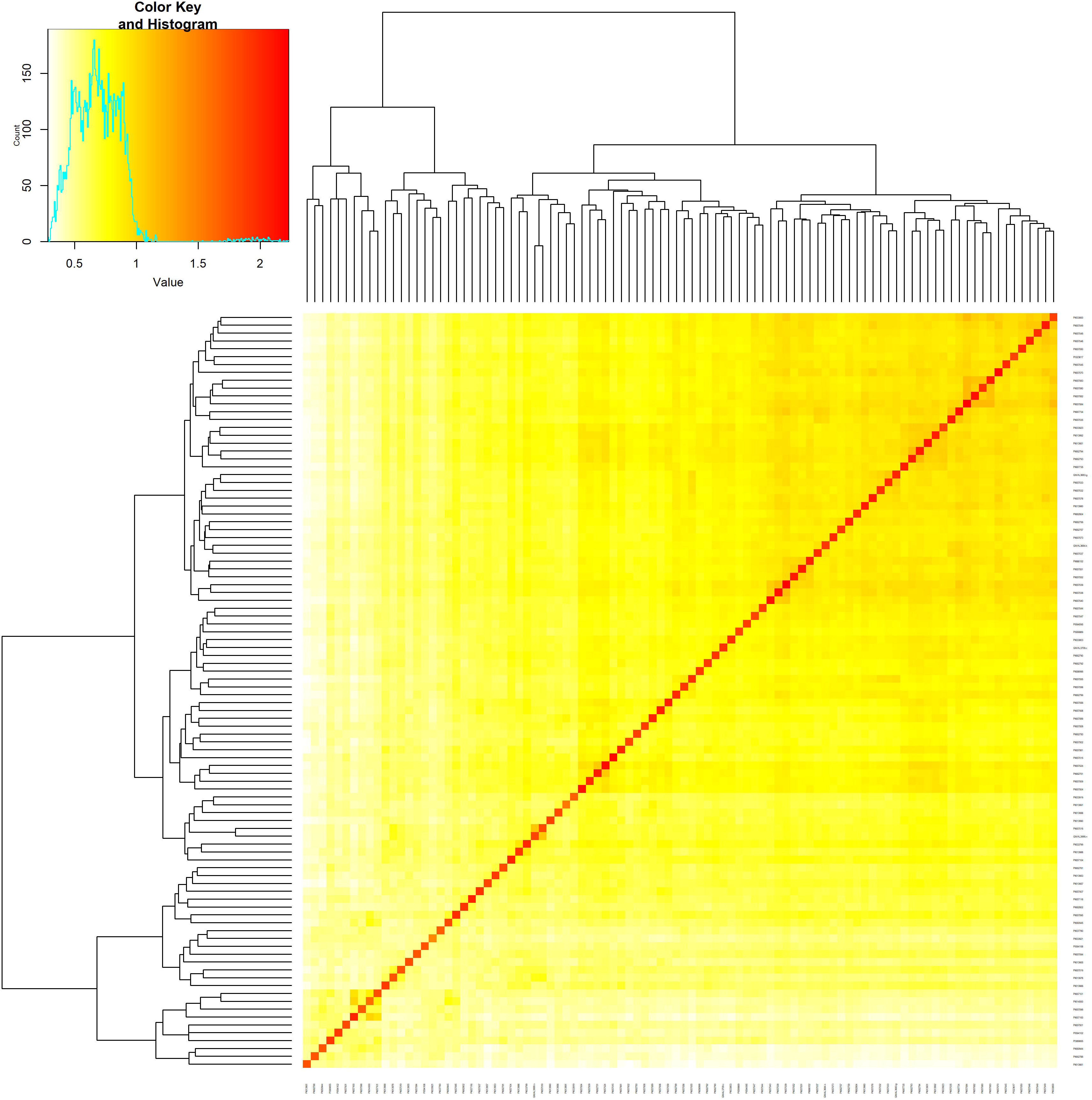
Kinship matrix of 96 *Malus* accessions used in GWAS.

**SFigure 2.**
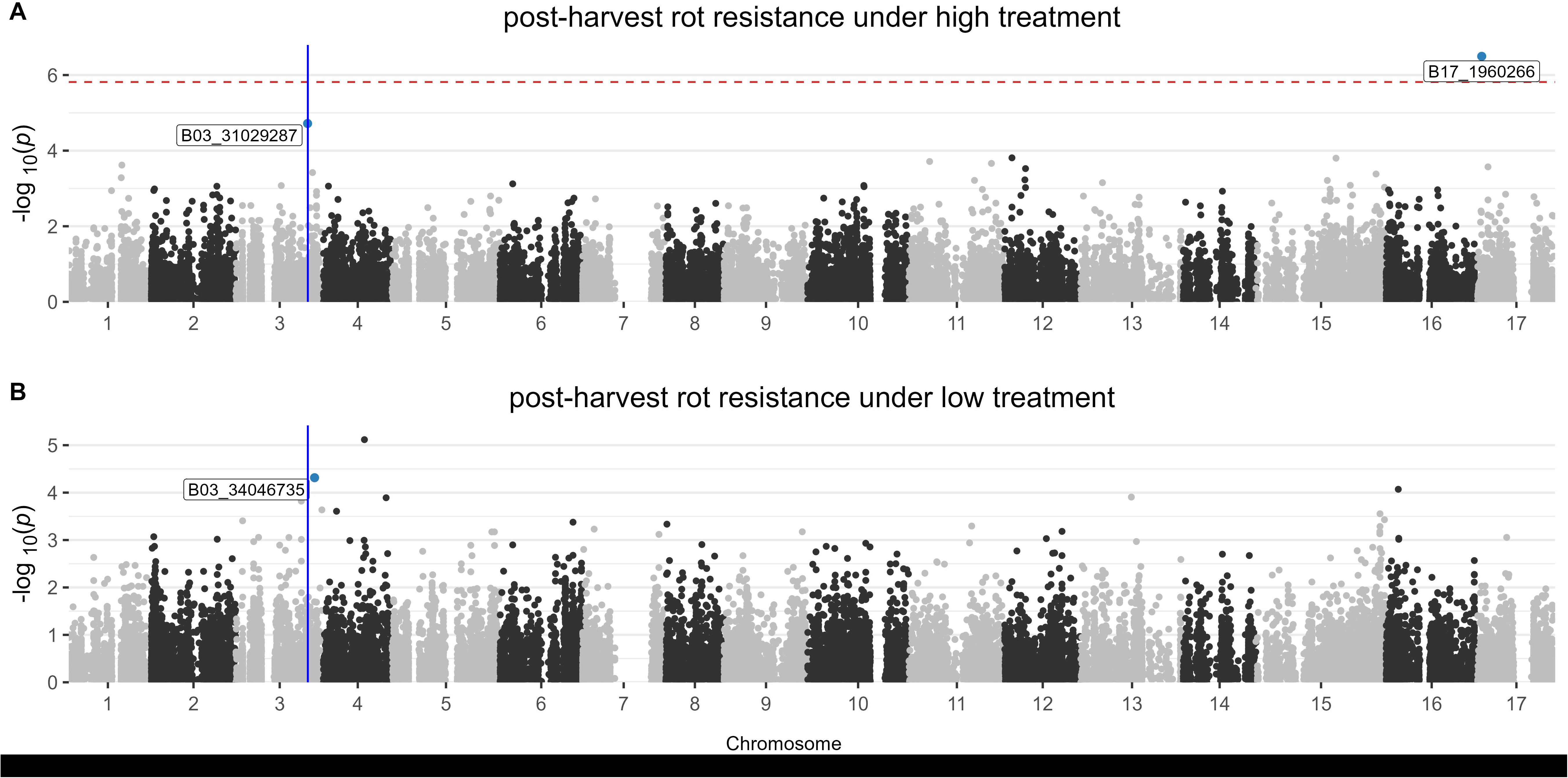
Manhattan’s of binary traits. Blue line represents the location of the *qM-Pe3.1* marker on chromosome 3.

**SFigure 3.**
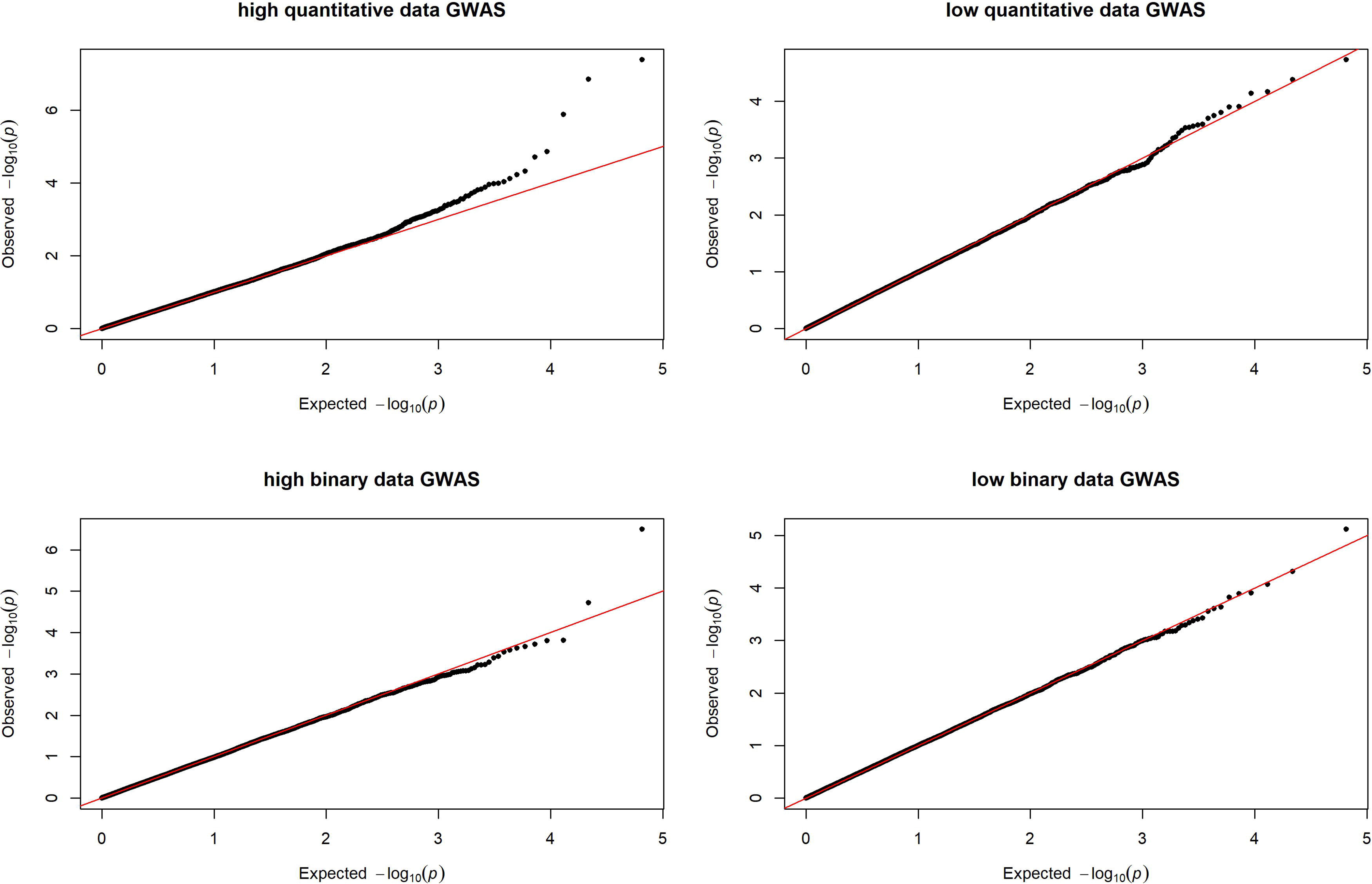
Quantile-quantile plots of each GWAS.

**SFigure 4.**
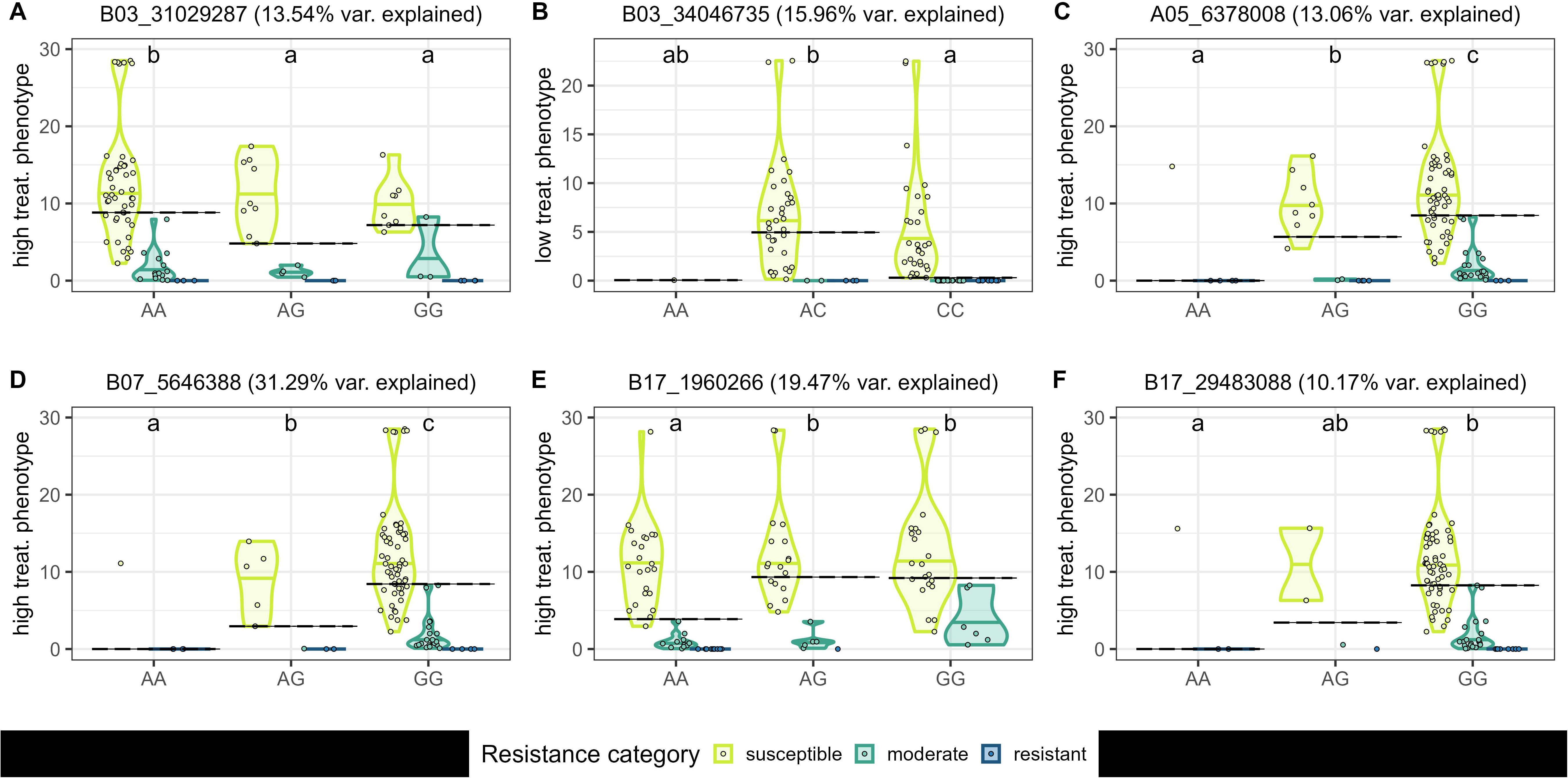
Distributions of phenotypes by allele and resistance category. Allelic effects of SNPs significantly associated with post-harvest rot in MLMM GWAS on: Chromosome 3 in the **A)** high binary data and **B)** low quantitative data (SNP B03_34046735 is also a cofactor in the low treatment binary data and explained 14.4% of phenotypic variance), **C)** Chromosome 5 in the high quantitative data, **D)** Chromosome 7 in the high quantitative data, and Chromosome 17 in the **E)** high binary data and **F)** high quantitative data. Points represent the mean lesion diameter of each accession. Accessions are grouped and colored by their *P. expansum* resistance category. Dashed bars represent the average across all resistance categories. Alleles not sharing any letter labeled at the top of the plot were determined significantly different by the Sidak-test (alpha 0.05).

